# A role for Toll-like receptor 3 in lung vascular remodeling associated with SARS-CoV-2 infection

**DOI:** 10.1101/2023.01.25.524586

**Authors:** Daniela Farkas, Srimathi Bogamuwa, Bryce Piper, Geoffrey Newcomb, Pranav Gunturu, Joseph S. Bednash, James D. Londino, Ajit Elhance, Richard Nho, Oscar Rosas Mejia, Jacob S. Yount, Jeffrey C. Horowitz, Elena A. Goncharova, Rama K. Mallampalli, Richard T. Robinson, Laszlo Farkas

## Abstract

Cardiovascular sequelae of severe acute respiratory syndrome (SARS) coronavirus-2 (CoV-2) disease 2019 (COVID-19) contribute to the complications of the disease. One potential complication is lung vascular remodeling, but the exact cause is still unknown. We hypothesized that endothelial TLR3 insufficiency contributes to lung vascular remodeling induced by SARS-CoV-2. In the lungs of COVID-19 patients and SARS-CoV-2 infected Syrian hamsters, we discovered thickening of the pulmonary artery media and microvascular rarefaction, which were associated with decreased TLR3 expression in lung tissue and pulmonary artery endothelial cells (ECs). *In vitro*, SARS-CoV-2 infection reduced endothelial TLR3 expression. Following infection with mouse-adapted (MA) SARS-CoV-2, TLR3 knockout mice displayed heightened pulmonary artery remodeling and endothelial apoptosis. Treatment with the TLR3 agonist polyinosinic:polycytidylic acid reduced lung tissue damage, lung vascular remodeling, and endothelial apoptosis associated with MA SARS-CoV-2 infection. In conclusion, repression of endothelial TLR3 is a potential mechanism of SARS-CoV-2 infection associated lung vascular remodeling and enhancing TLR3 signaling is a potential strategy for treatment.

## INTRODUCTION

Severe acute respiratory syndrome (SARS) coronavirus-2 (CoV-2) disease 2019 (COVID-19) is a deadly lung disease caused by SARS-CoV-2 ^1^. New variants continue to emerge, putting the health care system under strain ^2^. Long-term COVID-19 sequelae threaten the health and quality of life of COVID-19 survivors ^3^. Indeed, 25% of COVID-19 patients have long-term functional impairment and more than 70% have long-term radiological abnormalities in their lungs ^3^. Cardiovascular problems, such as myocarditis, heart failure, and disseminated microthrombosis, were discovered in COVID-19 patients ^4,5^. COVID-19 has been linked to an increased risk of right heart failure and remodeling ^6,7^. Furthermore, evidence suggests that COVID-19 patients are more likely to develop EC dysfunction, de-novo or worsening lung vascular remodeling, and pulmonary hypertension (PH) ^6–11^. COVID-19 causes more severe lung vascular remodeling than influenza ^9^. Lung vascular remodeling involves thickening of the pulmonary artery (PA) wall in COVID-19 patients, and COVID-19 patients have microvascular dysfunction ^9,11,12^. These changes are accompanied by decreased tissue perfusion, and microvascular distortion may be a trigger for intussusceptive lung angiogenesis ^12,13^.

In animal studies, endothelial cell (EC) apoptosis and dysfunction induce lung vascular remodeling and PH ^14,15^. COVID-19 not only causes morphological changes in ECs such as swelling and disruption of intercellular junctions ^13,16^, but also EC dysfunction as evidenced by EC barrier dysfunction, vascular inflammation, and thrombosis ^13,16^. Impaired EC function predicts a poor outcome in COVID-19 patients, and persistent EC dysfunction is suggested as one potential cause of long COVID-19 ^17,18^. As a result, ECs are expected to play a key role in COVID-19 pathogenesis. Inflammation, hypoxia, and direct virus infection are among the processes under study for COVID-19-associated EC dysfunction ^19^. Although there is some debate on the efficacy of SARS-CoV-2 to infect ECs due to the expression levels of the primary SARS-CoV-2 receptors in lung ECs, multiple reports describe evidence for direct infection of ECs by SARS-CoV-2 and alternative coreceptors that can facilitate entry of SARS-CoV-2 in ECs despite low expression of the primary receptor ^13,16,20–27^. However, the extent and mechanism of EC dysfunction and lung vascular remodeling after SARS-CoV-2 infection remain unknown.

Our group has recently discovered that deficiency of the innate immune receptor Toll-like receptor 3 (TLR3) promotes EC apoptosis, pulmonary artery remodeling, and PH ^28^. As an endosomal receptor for most ribonucleic acids (RNAs), including double stranded RNA and messenger RNA (mRNA), TLR3 plays an important role in the antiviral response ^29^. A recent study found that inherited TLR3 impairment amplifies SARS-CoV-2 replication *in vitro* and coincides with severe lung disease in COVID-19 patients ^30^. We have further shown that the TLR3 agonist Polyinosinic:polycytidylic acid [Poly(I:C)] reduces established PH in animal models without increasing lung vascular inflammation ^28^. We hypothesized that SARS-CoV-2 infection promotes lung microvascular and pulmonary artery remodeling and EC apoptosis *via* endothelial loss of TLR3. We further posit that treatment with Poly(I:C) protects from EC dysfunction and lung vascular remodeling caused by SARS-CoV-2 infection. In COVID-19 patients and SARS-CoV-2 infected Syrian hamsters, we discovered pulmonary artery wall thickening and a drop in microvascular density (MVD). TLR3 expression was reduced in COVID-19 patients and SARS-CoV-2 infected hamsters, as well as *in vitro* in SARS-CoV-2 infected human lung microvascular ECs (HLMVECs) and pulmonary artery ECs (PAECs). Finally, TLR3^-/-^ mice displayed increased pulmonary artery remodeling, whereas Poly(I:C) therapy protected from pulmonary artery remodeling and dysfunction, as well as microvascular pruning caused by infection with mouse-adapted (MA) SARS-CoV-2. Some of the data have been previously reported as a conference abstract ^31^.

## METHODS

### Reagents, cells, tissues, and constructs

#### Human cells and tissue

PAECs were obtained from the Pulmonary Hypertension Breakthrough Initiative (PHBI) and from the University of California Davis. Collection of PAECs without patient identifiers was approved at both institutions as non-human subjects research. Utilization of de-identified PAECs was deemed as non-human subjects research by the Office of Research Subjects Protection at OSU. Human lung microvascular ECs (HLMVECs) were purchased from Lonza clonetics. Human COVID-19 patient and control lung tissue samples were obtained from explanted lungs under Institutional Research Ethics Board approval # 2020H0306 (OSU). Informed consent and HIPAA authorization were obtained for patient samples. Associated clinical data were coded with a sample ID. Uncoded data were not shared with the investigators undertaking the study. Tissues were snap-frozen for RNA isolation or inflated with 10% Neutral Buffered Formalin, formalin-fixed, processed and paraffin-embedded. 5 μm sections were prepared. Control formalin-fixed, paraffin-embedded lung tissue sections were from failed donors and were obtained as de-identified tissue samples from the PHBI. These tissues were deemed non-human subjects research at OSU.

#### Antibodies

β-actin (A5441, MilliporeSigma, Saint Louis, MO, loading control), SARS-CoV-2 N protein (MA1-7404, Invitrogen), TLR3 (ab62566, Abcam, Cambridge, UK), TLR3 (6961, Cell Signaling Technology, Danvers, MA).

#### Kits and Reagents

For Terminal deoxynucleotidyl transferase dUTP nick end labeling (TUNEL), the Apoptag Fluorescein In Situ Apoptosis Detection Kit was used (S7110, MilliporeSigma). Texas Red-labeled Tomatolectin (*Lycopersicon esculentum* lectin, LEL) was purchased from Vector Laboratories, Newark, CA (TL-1176-1). 4’,6-Diamidino-2-phenylindole (DAPI) was purchased from MilliporeSigma (D9542). Endothelial growth media was purchased from Lonza Clonetics, Walkersville, MD (EGM-2MV, CC-3202). QIAzol was obtained from Qiagen, Germantown, MD (79306). Mayer’s Hematoxylin was purchased from Newcomer Supply (Middleton, WI).

#### Virus

SARS-CoV-2 (USA-WA1/2020, Batch # 70034262) and mouse-adapted SARS-CoV-2 (variant MA10, Cat # NR-55329) were obtained from Biodefense and Emerging Infections Research Resources (BEI Resources, Manassas, VA).

#### Animals

Syrian hamsters (*Mesocricetus auratus*) were obtained from Envigo (Indianapolis, IN, HsdHan:AURA). Wildtype (strain 101045) and TLR3^-/-^ mice (strain 005217) were obtained from Jackson Laboratories (Ellsworth, ME). The Syrian hamster tissues used in this study have been reported in a previous study ^32^.

#### Primers

SARS-CoV-2 Nuclecapsid (forward: CAATGCTGCAATCGTGCTAC; reverse: GTTGCGACTACGTGATGAGG; Integrated DNA Technologies, IDT), SARS-CoV-2 Envelope (forward: GGAAGAGACAGGTACGTTAA; reverse: AAGGTTTTACAAGACTCACG; IDT), SARS-CoV-2 Spike (forward: GCTGGTGCTGCAGCTTATTA; reverse: AGGGTCAAGTGCACAGTCTA; IDT), human *TLR3* (forward: FH2-TLR3; CAACAGAATCATGAGACAGAC; reverse: RH2-TLR3; CACTGTTATGTTTGTGGGTAG; Millipore Sigma), *GUSB* (forward: FH2-GUSB; CTCCAGCTTCAATGACATC; reverse: RH2-GUSB; ATTCACCCACACGATGG, Millipore Sigma), mouse *Il6* (forward: TACCACTTCACAAGTCGGA; reverse: AATTGCCATTGCACAACTC; IDT), mouse *Il1b* (forward: CCTCAAAGGAAAGAATCTATACCTG; reverse: CTTGGGATCCACACTCTCC, IDT), mouse *Tnfa* (forward: TTCTCATTCCTGCTTGTGG; reverse: TTGGGAACTTCTCATCCCT; IDT), mouse *Cxcl10* (forward: GACTCAAGGGATCCCTCTC; reverse: ATGGCCCTCATTCTCACTG; IDT), mouse *Il10* (forward: TTAATAAGCTCCAAGACCAAGG; reverse: CATCATGTATGCTTCTATGCAG; IDT), mouse *Gusb* (forward: ATTCAGATATCCGAGGGAAAG, reverse: TCCTCTGAGTAGGGATAGTG; IDT), mouse *Tbp* (forward: GTTCTTAGACTTCAAGATCCAG; reverse: TTCTGGGTTTGATCATTCTG; IDT).

### Animal experiments

Snap-frozen and formalin-fixed, paraffin-embedded lung and heart tissues from SARS-CoV-2 infected and control Syrian Hamsters (*Mesocricetus auratus*) were obtained from a previously published study ^32^. All biosafety level 3 (BSL3) experiments were performed in the OSU BSL3 facility in adherence to approved biosafety protocols. For mouse experiments, male mice received MA SARS-CoV-2 or mock infection (PBS) as published in our previous study ^33^. In brief, mice were anesthetized with isoflurane, weighed, and held in a semi-supine position. 50 μL of MA SARS-CoV-2-containing PBS (2.5×10^4^ PFU) or PBS alone (control mice) was given intranasally. We have published the efficiency of MA SARS-CoV-2 infection recently ^33^. Some mice were treated with 10 mg/kg Poly(I:C) or vehicle three times a week by intraperitoneal injection, similar to our previous publication ^28^. Mice were randomly assigned to treatment or vehicle groups and body weight was regularly recorded. At day 7, mice underwent echocardiography with a Vivid iQ R3 Premium and a L8-18i-RS transducer (GE Healthcare, Chicago, IL) under ketamine/xylazine anesthesia. After echocardiography, blood was collected, the mice were euthanized by exsanguination and lung and heart tissue were harvested.

### Cell culture experiments

PAECs and HLMVECs were grown in complete EGM-2MV (Lonza). For SARS-CoV-2 infection, cells were infected with 2 MOI of SARS-CoV-2 and cells were removed in QIAzol after 6h for RNA isolation.

### Protein isolation and immunoblotting

Hamster lung tissue was homogenized, and protein lysate was prepared as published previously ^32^. SDS-PAGE and immunoblotting were performed as published previously ^28,32,34^. The immunoblots were imaged using the ChemiDoc chemiluminescence imaging system (BioRad). Images were analyzed for densitometry with automated unbiased background subtraction using ImageLab 6.1 (BioRad).

### RNA isolation and real-time quantitative PCR

Total RNA was isolated from cells and lung tissue as previously published using Qiagen RNeasy kits ^28,34^. Cells were scraped in QIAzol and lung tissue was homogenized in QIAzol in inactive SARS-CoV-2 prior to removal of tissue from the BSL3 facility as approved by the Institutional Biosafety Committee. Reverse transcription and quantitative real-time PCR was performed according to a previously published protocol ^28,34^ with a BioRad CFX384 system.

### Histology and immunohistochemistry

Formalin-fixed and paraffin-embedded tissue was stained with hematoxylin-eosin (H&E) or Masson Trichrome (Histowiz, Brooklyn, NY) or stained according to established immunohistochemistry (IHC) protocols for TLR3 and cleaved caspase-3 ^28^. H&E and Masson Trichrome images were automatically acquired by Histowiz. IHC images were acquired with an EVOS M7000 automated microscope (ThermoFisher Scientific). The ApopTag Peroxidase In Situ Apoptosis Detection Kit was used for Terminal deoxynucleotidyl transferase dUTP nick end labeling (TUNEL) (Sigma Millipore) according to the manufacturer’s instructions, followed by counterstaining with Mayer’s Hematoxylin. Images were taken with a EVOS M7000 automated microscope. To identify SARS-CoV-2-infected ECs, ECs were labeled with Texas Red-conjugated Tomatolectin (*Lycopersicon esculentum*, LEL) and stained with anti-SARS-CoV-2 N protein antibody similar to previously published protocols ^34^. Images were taken with an EVOS M7000 automated microscope and an Olympus FV3000 confocal microscope (Campus Microscopy and Imaging Facility, OSU). Media wall thickness (MWT) was measured and calculated as published previously ^28,34–36^. In brief, H&E stained sections were prepared and whole slides were scanned by Histowiz. Images containing pulmonary arteries were downloaded and external diameter (ED) and internal diameter (ID) were measured using ImageJ. Media thickness (MT) was calculated as 2×MT=ED-ID. MWT was calculated as *MWT* = (2 X *MT/ED*) X 100%. Pulmonary arteries were differentiated into small-sized 25 μm < ED < 50 m and medium-sized 50 μm ≤ ED < 100 μm for hamsters. In mice, pulmonary arteries size 25-100 μm were included. The number of N^+^ LEL^+^ cells and LEL^+^ cells was counted per field of view in 10 images/animal. The fraction of N^+^ LEL^+^ vs. all LEL^+^ cells was calculated. For calculating microvascular density (MVD), tissue sections were stained with LEL and DAPI.

Images were taken with an EVOS M7000 automated microscope. The microvascular area was measured as Texas Red^+^ LEL staining vs. total tissue area in 10 images/animal. For detection of microvascular apoptosis, sections were stained for TUNEL with the ApopTag Fluorescein In Situ Apoptosis Detection Kit (Sigma Millipore) followed by EC staining with Texas Red-conjugated Tomatolectin. Z-stacks were acquired with an Olympus FV3000 confocal microscope (Campus Microscopy and Imaging Facility, OSU). The nucleus in fluorescence stainings was stained with DAPI. For measurement of RV CM CSA, Masson Trichrome stained slides were prepared by Histowiz and whole slides were scanned by Histowiz. Random images were downloaded from Histowiz. CM CSA was measured using Fiji by an investigator blinded to the treatment groups from at least 4 separate areas per animal. Blinded analysis in all histological quantification was ensured by numerical coding of treatment groups.

### Statistical analysis

Normal distribution of data was tested by the Shapiro-Wilk test. Normally distributed data were compared using Student’s t-test (two groups), or one or two-way ANOVA (more than two groups), followed by multiple comparison correction using the Holm-Sidak or Sidak tests. Nonparametric data were compared by the Mann-Whitney U test (two groups) or Kruskal-Wallis analysis followed by multiple comparison correction according to Dunn. The calculations were performed using Prism 9 (GraphPad Software Inc., San Diego, CA). A P value of <0.05 was considered significant.

## RESULTS

### SARS-CoV-2 infection induces pulmonary artery and lung microvascular remodeling and apoptosis and loss of TLR3 expression

To determine if SARS-CoV-2 infection causes lung vascular remodeling and apoptosis, we performed morphometric analysis on lung tissue sections from human control patients and COVID-19 patients, as well as lung tissue from SARS-CoV-2 infected Syrian hamsters. We detected persistent thickening of the pulmonary artery wall by media wall thickness (MWT) and loss of lung tissue microvascular density (MVD) in human COVID-19 patients (**Fig. 1A-D**). In addition, TLR3 expression was reduced in PAECs and in lung tissue from COVID-19 patients (**Fig. 1E-F**). To identify if endothelial infection with SARS-CoV-2 directly decreases TLR3 expression, we infected human PAECs and HLMVECs with SARS-CoV-2 and measured SARS-CoV-2 nucleocapsid (N), envelope (E), and spike (S) protein mRNA to confirm EC infection, as well as TLR3 mRNA expression. We detected N, S, and E protein mRNA after 6h of infection, and TLR3 was reduced after 6h in PAECs and HLMVECs (**Fig. 1G-J**). In a similar manner, SARS-CoV-2 infection induced persistent pulmonary artery media thickening and reduced MVD in Syrian hamsters (**Fig. 2A-E**). In the lungs of SARS-CoV-2 infected hamsters, we also found microvascular ECs expressing N protein, indicating *in vivo* infection of ECs (**Fig. 2F-G**). Testing a potential mechanism for lung vascular remodeling, we also detected more apoptotic ECs in pulmonary arteries and alveolar walls (**Fig. 2H-J**). These findings were associated with reduced expression of TLR3 in PAECs *in situ* and in lung tissue following SARS-CoV-2 infection (**Fig. 2I, K**). Supporting our findings of lung vascular remodeling, we detected increased right ventricular (RV) cardiomyocyte (CM) cross-sectional area (CSA) in the RVs of SARS-CoV-2 infected hamsters (**Fig. 2L-M**).

**Figure 1.**
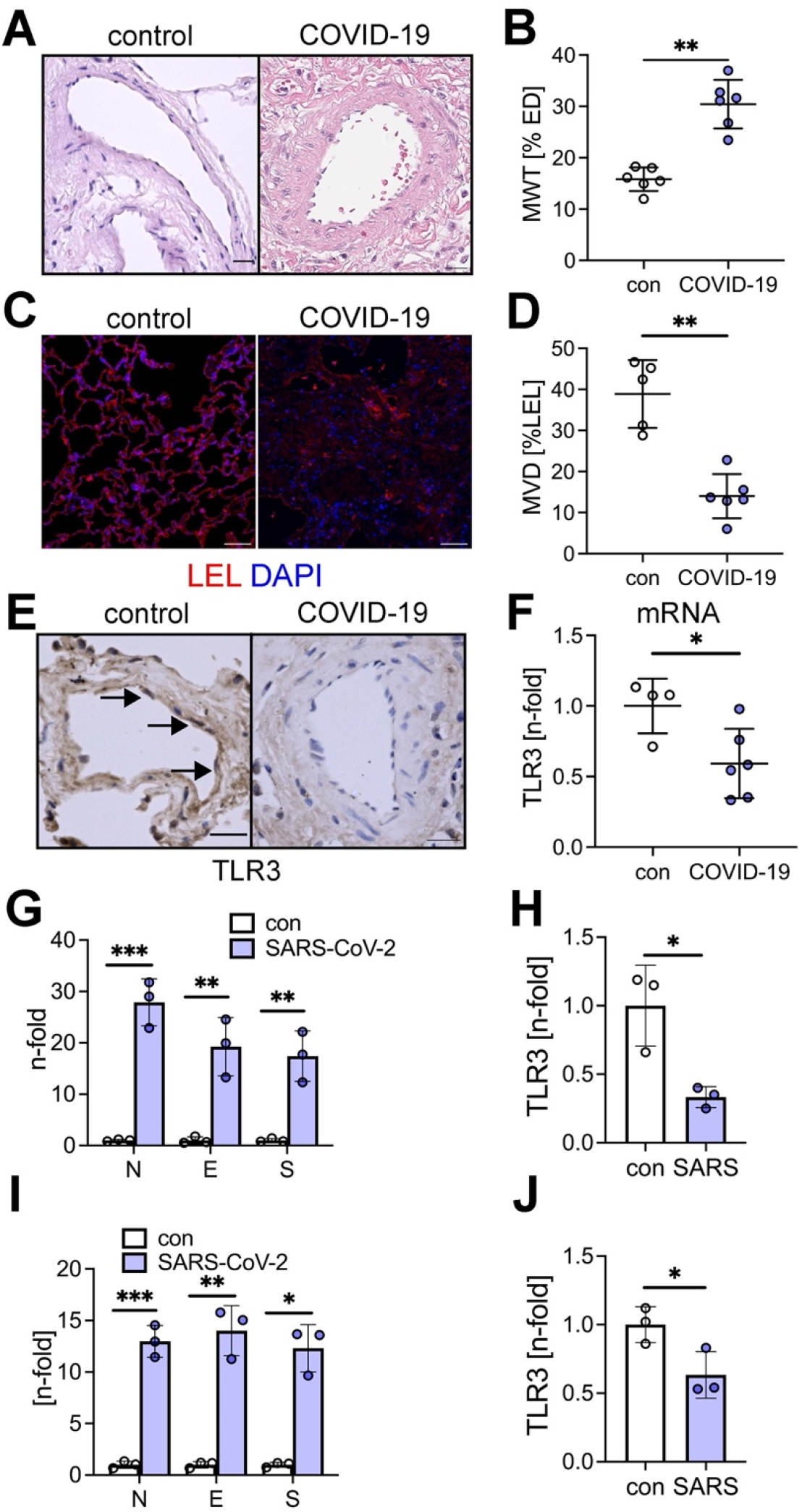
SARS-CoV-2 infection induces lung vascular remodeling and impairs TLR3 expression in human patients. (**A**) Representative Hematoxylin and Eosin (H&E) sections of human control and COVID-19 lungs (explant tissue) show thickening of the pulmonary artery wall following SARS-CoV-2 infection. (**B**) Quantification of media wall thickness (MWT) of pulmonary arteries in control and COVID-19 patients. (**C**) Comparison of microvascular density (MVD) using Tomatolectin staining (*Lycopersicon esculentum*, LEL) in control and COVID-19 patients (explant lung tissue). (**D**) Quantification of MVD as % LEL staining. (**E**) Representative images of pulmonary arteries stained for TLR3 immunohistochemistry (IHC) demonstrate reduced endothelial TLR3 expression in COVID-19 patients (day 4). Arrows in indicate strong TLR3 staining in PAECs from control patient. (**F**) qRT-PCR of lung tissue from control and COVID-19 patients demonstrates reduced lung tissue mRNA expression of TLR3 in COVID-19 lungs. (**G-J**) Infection of human PAECs (G, H) and HLMVECs (I, J) with SARS-CoV-2 (MOI 2) shows induction of mRNA for SARS-CoV-2 nucleocapsid (N), envelope (E), and spike (S) proteins in PAECs (G) and HLMVECs (I) after 6h, indicating virus replication. Note that the C_T_ value of controls was measured at the level of RT^-^ and H_2_O controls. SARS-CoV-2 infection reduced mRNA expression of TLR3 in PAECs (H) and HLMVECs (J) after 6h. Scale bars: 25μm (E), 50μm (A), 100μm (C). Nuclear staining with DAPI (C). Counterstaining with Mayer’s Hematoxylin (E). Data are presented as single values and mean±SD. *P<0.05, **P<0.01, ***P<0.001.

**Figure 2.**
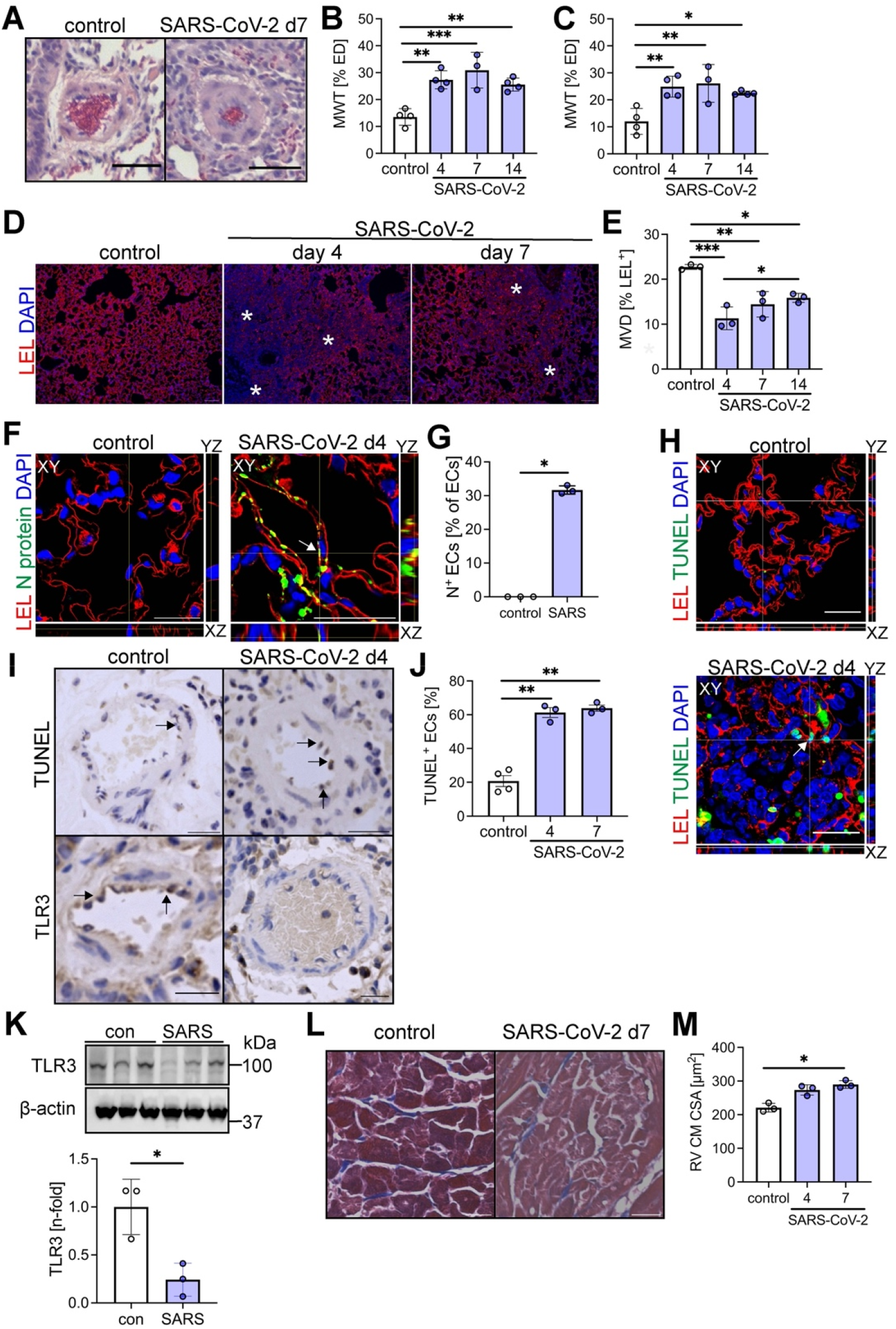
SARS-CoV-2 infection induces lung vascular remodeling, endothelial apoptosis, and represses TLR3 expression in Syrian hamsters. (**A**) Representative H&E sections of PBS and SARS-CoV-2 treated hamsters (day 7) show thickening of the pulmonary artery wall following SARS-CoV-2 infection. Control animals received PBS. (**B-C**) Quantification of media wall thickness (MWT) of small (B, 25μm<external diameter, ED<50μm) and medium (C, 50μm<ED≤100μm) pulmonary arteries in SARS-CoV-2 infected hamsters (days 4, 7 and 14) and controls. (**D**) Representative images show loss of MVD in LEL-stained lung tissue sections from control and SARS-CoV-2 infected hamsters (days 4, 7). Asterisks indicate areas of reduced MVD. (**E**) Quantification of MVD as % LEL staining area vs. total tissue area. (**F**) Representative orthogonal sections from z-stacks (confocal microscopy) show SARS-CoV-2 N protein expression in lung ECs (representative cell labeled with arrow) at day 4 following SARS-CoV-2 infection. (**G**) Quantification of SARS-CoV-2 N protein expression in lung ECs (arrows) at day 4. (**H**) Representative orthogonal sections from z-stacks (confocal microscopy) show TUNEL^+^ microvascular ECs (LEL^+^) in SARS-CoV-2 infected hamsters at day 4 (arrow), but not in control hamsters. (**I**) Representative images of TUNEL staining indicates EC apoptosis (arrows) in pulmonary arteries of SARS-CoV-2 infected hamsters. Representative images of TLR3 IHC show loss of endothelial TLR3 expression in pulmonary arteries from SARS-CoV-2 infected hamsters at day 4. ECs in pulmonary arteries of control animals have strong TLR3 staining (arrows). (**J**) Quantification of TUNEL^+^ ECs in pulmonary arteries of SARS-CoV-2 infected hamsters. (**K**) Reduced TLR3 expression in the lung tissue protein lysate of control and SARS-CoV-2 infected hamsters at day 4. The figures show the immunoblot of TLR3 and loading control β-actin, and densitometric quantification after automated background adjustment. Data are expressed as n-fold of control. (**L-M**) Representative Masson Trichrome images (L) and quantification of right ventricular (RV) cardiomyocyte (CM) cross-sectional area (CSA, M) indicate increase in RV CM CSA following SARS-CoV-2 infection. Scale bars: 20 μm (I, TLR3 staining), 25μm [A, F, H, I (TUNEL), L), 100μm (D). Nuclear staining with DAPI (D, F, and I). Data in graphs are presented as single values and mean±SD. *P<0.05, **P<0.01, ***P<0.001.

### The TLR3 agonist Poly(I:C) protects and TLR3 knockout exaggerates SARS-CoV-2 induced lung vascular remodeling

To test if manipulation of TLR3 signaling and expression modulates the severity of SARS-CoV-2 induced lung vascular remodeling, we infected mice with MA SARS-CoV-2 and treated those mice with the TLR3 agonist Poly(I:C). In addition, we compared TLR3^+/+^ and TLR3^-/-^ mice following MA SARS-CoV-2 infection. We found substantial inflammation and transient weight loss with MA SARS-CoV-2 infection in TLR3^+/+^ mice and reduced lung inflammation and lesser weight loss with Poly(I:C) treatment and TLR3 knockout following MA SARS-CoV-2 infection (**Fig. 3A-C**). Further, we found perivascular inflammation and increased MWT in MA SARS-CoV-2 infected TLR3^+/+^ mice, which was prevented by Poly(I:C) treatment. In contrast, TLR3^-/-^ mice had exaggerated MWT following MA SARS-CoV-2 infection (**Fig. 3D-E**). Likewise, MA SARS-CoV-2 infection reduced MVD in TLR3^+/+^ and TLR3^-/-^ mice and Poly(I:C) treatment prevented the loss of MVD following MA SARS-CoV-2 infection (**Fig. 3D, F**). However, TLR3^-/-^ mice showed no exaggerated loss of MVD compared to TLR3^+/+^ mice. We detected increased EC apoptosis in TLR3^+/+^ +SARS-CoV-2 mice, which was reduced by Poly(I:C) treatment and exaggerated in TLR3^-/-^ mice (**Fig. 3D, G**). Testing RV remodeling, we found that SARS-CoV-2 infection increased RV CM CSA, and this effect was similar in TLR3^+/+^ SARS-CoV-2+veh, TLR3^+/+^ SARS-CoV-2+Poly(I:C), and TLR3^-/-^ SARS-CoV-2 treated mice (**Fig. 3D, H**). We further tested the effect of SARS-CoV-2 infection on the echocardiographic parameters of cardiac function. We detected a reduced ratio of pulmonary artery acceleration time (PAT) vs. pulmonary ejection time (PET), an inverse indicator of pulmonary artery pressure, in MA SARS-CoV-2 infected TLR3^+/+^ mice, which was prevented by Poly(I:C) treatment. TLR3^-/-^ mice showed a mild exaggeration of PAT/PET reduction following MA SARS-CoV-2 infection compared to TLR3^+/+^ mice (**Fig. 3I**). Left ventricular ejection fraction (LV EF) was reduced in TLR3^-/-^ mice following SARS-CoV-2 infection, but not in TLR3^+/+^ +SARS-CoV-2 mice, indicating the protective role of TLR3 in the LV during virus infection (**Fig. 3J**).

**Figure 3.**
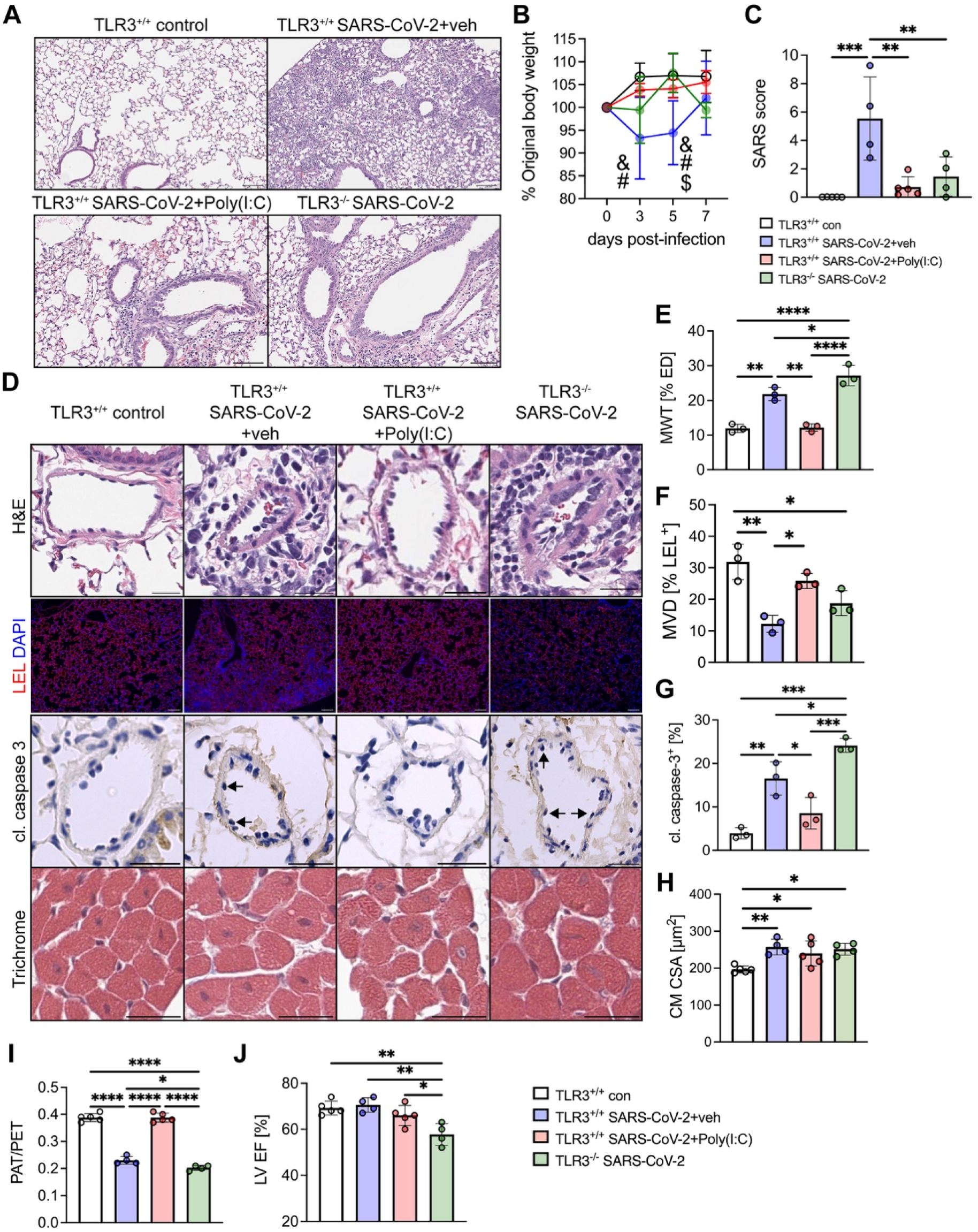
TLR3 agonist Poly(I:C) protects from lung vascular remodeling, whereas TLR3 knockout exaggerates lung vascular remodeling following infection with mouse-adapted SARS-CoV-2. (**A**) Representative low power images (H&E) show substantial tissue inflammation seven days after infection with mouse-adapted (MA) SARS-CoV-2. (**B**) Weight curves of the four groups are shown as % of original body weight. & P<0.001 TLR3^+/+^ con vs. TLR3^+/+^ SARS-CoV-2+veh; # P<0.01 TLR3^+/+^ SARS-CoV-2+veh vs. TLR3^+/+^ SARS-CoV-2+Poly(I:C) (10 mg/ml by intraperitoneal injection, three times a week); $ P<0.001 TLR3^+/+^ SARS-CoV-2+veh vs. TLR3^-/-^ SARS-CoV-2 (ANOVA). (**C**) Average SARS score of the four groups. (**D**) Representative H&E images of pulmonary arteries, representative LEL immunofluorescence indicating microvascular density, representative cleaved caspase-3 IHC and representative images of Masson Trichrome stained right ventricles. (**E**) Quantification of MWT. (**F**) Quantification of MVD. (**G**) Quantification of cleaved caspase-3^+^ ECs in PAs. (**H**) Quantification of right ventricular (RV) cardiomyocyte cross-sectional area (CM CSA). (**I**) ratio of pulmonary artery acceleration time (PAT) and pulmonary ejection time (PET) obtained by pulse-wave doppler echocardiography. (**J**) Quantification of left ventricular ejection fraction (LV EF) by echocardiography and estimation according to Teich. Scale bars: 25μm (D, images with H&E, cleaved caspase-3, and Trichrome staining), 100μm (D, images with LEL staining), 200μm (A). Data in graphs are presented as single values and mean±SD. *P<0.05, **P<0.01, ***P<0.001, ****P<0.0001.

### TLR3 regulates expression of SARS-CoV-2proteins and inflammatory cytokines

MA SARS-CoV-2 infected mice showed induction of SARS-CoV-2 N, S, and E protein mRNA, which was reduced with Poly(I:C) treatment at a trend. While TLR3^-/-^ mice had reduced expression of SARS-CoV-2 N and S proteins, E protein expression remained high in TLR3^-/-^ mice (**Fig. 4**). MA SARS-CoV-2 infection caused a trend towards *Il6, Il1b, Tnfa, Cxcl10* and *Il10*, similar to previous description in TLR3^+/+^ mice ^37^. Poly(I:C) amplified the expression of these cytokines and chemokines, showing the induction of protective immune signaling (**Fig. 4**). TLR^-/-^ mice had a lower expression of these cytokines and chemokines following MA SARS-CoV-2 infection (**Fig. 4**).

**Figure 4.**
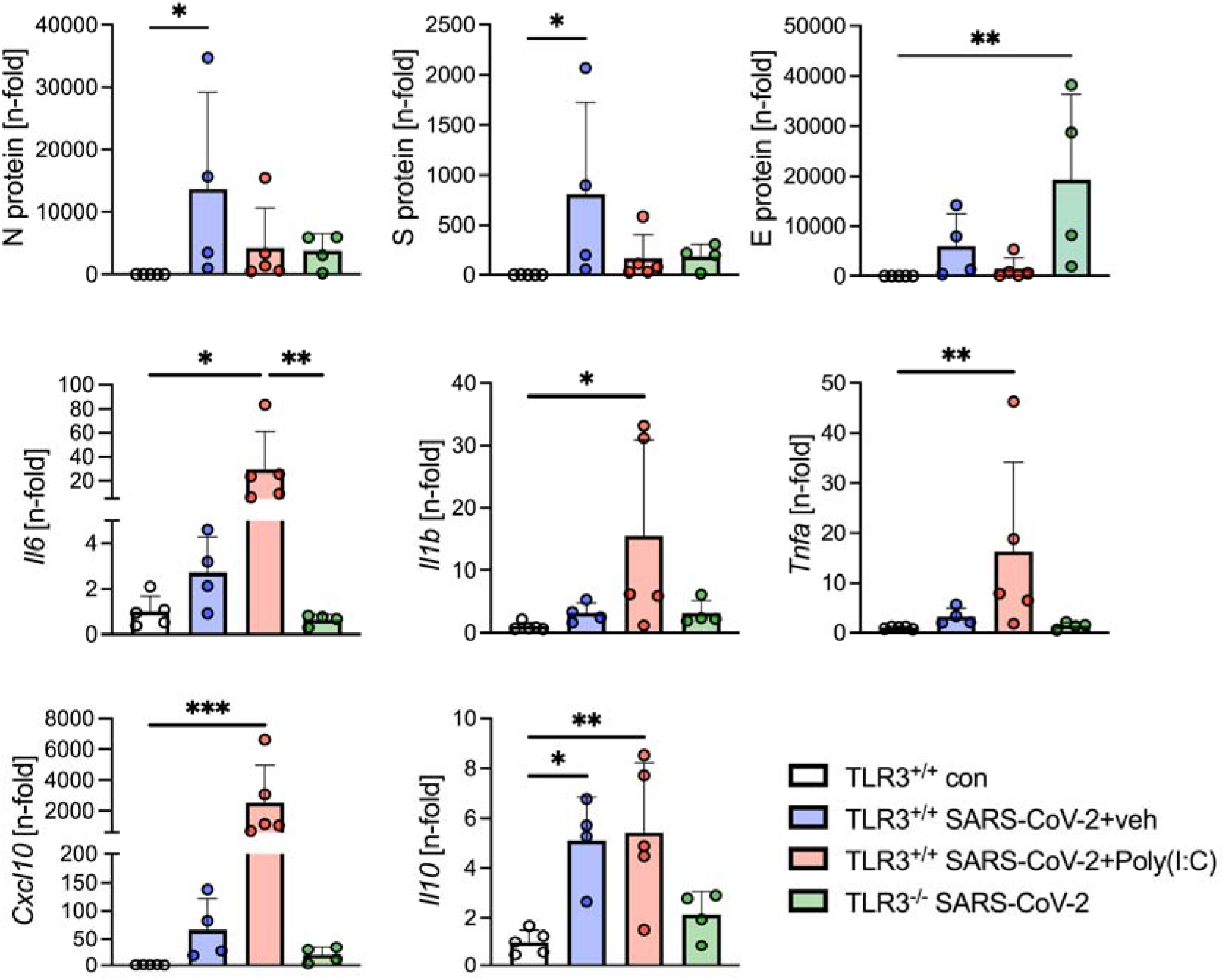
qRT-PCR of SARS-CoV-2 viral proteins and inflammatory marker profile in the lungs of MA SARS-CoV-2 infected mice. Diagrams show mRNA expression by qRT-PCR for SARS-CoV-2 N, S, and E proteins, as well as mouse *Il6, Il1b, Tnfa, Cxcl10, Il10*. The following groups are compared: TLR3^+/+^ control, TLR3^+/+^ +SARS-CoV-2+vehicle, TLR3^+/+^ +SARS-CoV-2+Poly(I:C) (10 mg/ml by intraperitoneal injection, three times a week) and TLR3^-/-^ +SARS-CoV-2. Data are presented as single values and mean+SD. \**P*<0.05, \*\**P*<0.01, \*\*\**P*<0.001.

## DISCUSSION

The cardiovascular complications of COVID-19 affect patients with acute and long COVID-19 ^4,5^. EC dysfunction, remodeling of the lung vasculature, including pulmonary artery wall thickening, and PH are emerging COVID-19 consequences ^6–9,11^. In this work we evaluated the idea that SARS-CoV-2 infection induces micro- and macrovascular lung vascular remodeling and EC apoptosis, and that endothelial TLR3 loss is an underlying mechanism.

The main findings of our study were as follows: a) lungs of patients with COVID-19 and of SARS-CoV-2-infected Syrian hamsters show pulmonary artery wall thickening, reduction of microvascular density (MVD), and reduced TLR3 expression; b) endothelial apoptosis is augmented in the lungs of SARS-CoV-2 infected hamsters; c) SARS-CoV-2 infection repressed TLR3 in HLMVECs and PAECs *in vitro*; d) TLR3^-/-^ mice had exaggerated pulmonary artery remodeling, whereas treatment with Poly(I:C) protected them from pulmonary artery remodeling, pulmonary artery dysfunction, and the microvascular pruning associated with infection with MA SARS-CoV-2.

Early in the COVID-19 pandemic data appeared indicating right ventricular strain and remodeling, as well as heterogenous pulmonary perfusion and pulmonary vascular remodeling, in patients with COVID-19 ^9,38,39^. Furthermore, reports showed the emergence of PH in patients who had no prior history of lung vascular disease ^7,9^. Our new systematic analysis demonstrates that COVID-19 patients receiving a lung transplant exhibited considerable thickening of the pulmonary artery media and significant capillary loss, which is consistent with destruction of the alveolar tissue architecture. To better understand the role of SARS-CoV-2 in this process, we used two models of the SARS-CoV-2 infection: The Syrian hamster model ^32,40,41^ and the MA SARS-CoV-2 murine infection model ^33,37,42^. In this study, we show that SARS-CoV-2 infection produces substantial microvascular and pulmonary artery remodeling in hamsters, and that this remodeling can be detected early on and persists beyond the acute viral replication, which we have characterized previously ^32^. While decreased perfusion and a heterogenous perfusion pattern have previously been observed in COVID-19 patients, earlier reports have concentrated on extensive thrombosis ^38,43–45^. SARS-CoV-2 causes lung microvascular endothelial apoptosis, which is one possible mechanism for microvascular dropout ^35,46,47^. Endothelial apoptosis is a known driver of pulmonary artery remodeling, and we found EC apoptosis in pulmonary arteries and increased pulmonary artery muscularization in the lungs of SARS-CoV-2 infected hamsters, and EC apoptosis is consistent with previous findings in lungs from human COVID-19 patients and non-human primates infected with SARS-CoV-2 ^35,46,47^.

We investigated how SARS-CoV-2 infection changes TLR3 expression *in vitro* and *in vivo* to better understand the probable mechanism underpinning SARS-CoV-2-induced lung vascular remodeling. TLR3 is an important component of the antiviral response because it recognizes viral RNAs in the endosome and induces a type I interferon response ^29^. SARS-CoV-2 infection lowers TLR3 expression in lung ECs *in vitro* and *in vivo*. Furthermore, our findings imply that whole body knockout of TLR3 increases SARS-CoV-2 induced pulmonary artery remodeling and further reduces PAT/PET. The latter indicates increased pulmonary artery pressure. TLR3 knockout also decreases LV function after SARS-CoV-2 infection but has no effect on MVD. We prevented SARS-CoV-2 induced pulmonary artery muscularization, PAT/PET reduction, and loss of MVD by using the TLR3 agonist Poly(I:C). These findings are consistent with prior observations in the systemic and pulmonary circulation by others and us, demonstrating that TLR3 deficiency makes ECs more vulnerable to apoptosis ^28,48^. As a result, augmented EC dysfunction caused by TLR3 knockout is a putative driver of pulmonary artery muscularization, explaining the exacerbated drop in PAT/PET. Furthermore, TLR3 protects against adverse cardiac remodeling and dysfunction in the context of various viral infections, which could explain the effect of TLR3 knockout on LV function in our study ^49–51^. However, TLR3 signaling could also be promoting vascular and cardiac remodeling in a model- or context-dependent manner, according to certain findings ^52–54^. While TLR3 agonist treatment generates protective immunity to minimize the harm caused by SARS-CoV-2 infection ^55,56^, our data are the first to show that TLR3 protects pulmonary circulation and the heart after SARS-CoV-2 infection.

TLR3^-/-^ mice also appear to have a gap between the degree of pulmonary artery remodeling and inflammation. One possible reason for this counterintuitive outcome might be the use of whole-body TLR3^-/-^ in our study; TLR3 is a plausible mediator of inflammation in epithelial cells after SARS-CoV-2 infection and may contribute to lung inflammation through the TLR3-mediated activation of immune cells ^57^. Furthermore, TLR3^-/-^ mice also show a discrepancy between the expression of SARS-CoV-2 proteins; while mRNA expression of SARS-CoV-2 E protein was very high in TLR3^-/-^ mice, N and S protein expression was reduced. In contrast, the TLR3 agonist Poly(I:C) induced protective inflammatory cytokine expression, including IL-10 and TNF-α ^56,58^, and reduced the inflammatory score and mRNA expression of SARS-CoV-2 proteins. These results support a protective, anti-viral effect in addition to increased microvascular density and reduced pulmonary artery MWT and EC apoptosis.

We acknowledge the following limitation of our study: because our treatment with Poly(I:C) was administered in a preventive manner, it is difficult to disentangle the effects on lung inflammation and pulmonary vascular remodeling following SARS-CoV-2 infection.

In conclusion, SARS-CoV-2 infection causes microvascular dropout and pulmonary artery remodeling associated with endothelial apoptosis. One potential mechanism is repression of TLR3 expression in ECs and enhancing TLR3 signaling may serve as an approach for reducing lung vascular remodeling associated with SARS-CoV-2 infection.

## ACKNOWLEDGEMENTS

The authors acknowledge the expert technical assistance of Jaylen Hudson. Specimens, and/or data were provided by The Ohio State University Wexner Medical Center Comprehensive Transplant Center Human Tissue Biorepository. Data/Tissue samples were further provided under the Pulmonary Hypertension Breakthrough Initiative (PHBI).

## SOURCES OF FUNDING

The study was supported by grants from the NIH/NHLBI (HL139881 to L.F., HL141195 to J.C.H., HL150638, HL120261 and HL113178 to E.A.G.) and institutional start-up funds from The Ohio State University to L.F. The study was further supported by Award Number Grant UL1TR002733 and KL2TR002734 from the National Center for Advancing Translational Sciences to J.S.B. Funding for the PHBI is provided under an NHLBI R24 grant, R24HL123767, and by the Cardiovascular Medical Research and Education Fund (CMREF). The authors acknowledge resources for confocal microscopy from the Campus Microscopy and Imaging Facility (CMIF), and the OSU Comprehensive Cancer Center (OSUCCC) Microscopy Shared Resource (MSR) at The Ohio State University with NIH S10 OD025008 and NIH NIC P30CA016058. These facilities are supported in part by grant P30 CA016058, National Cancer Institute, Bethesda, MD. The content is solely the responsibility of the authors and does not necessarily represent the official views of the National Institutes of Health.

## DISCLOSURES

The authors declare no competing financial interests.

## HIGHLIGHTS

- The lungs of patients with COVID-19 and of SARS-CoV-2-infected Syrian hamsters show pulmonary artery wall thickening, reduction of microvascular density, and reduced TLR3 expression.
- Endothelial apoptosis is augmented in the lungs of SARS-CoV-2 infected hamsters.
- SARS-CoV-2 infection repressed TLR3 in human lung microvascular endothelial cells and pulmonary artery endothelial cells *in vitro*
- TLR3^-/-^ mice had exaggerated pulmonary artery remodeling, whereas treatment with the TLR3 agonist polyinosinic:polycytidylic acid protected them from pulmonary artery remodeling, pulmonary artery dysfunction, and the microvascular pruning associated with infection with mouse-adapted SARS-CoV-2.

## Notes

### Competing Interest Statement

The authors have declared no competing interest.

